# Bayesian Estimation of Mosaic Loss of Chromosome Y from Bulk RNA Sequencing Data

**DOI:** 10.64898/2026.05.20.726153

**Authors:** Jhih-Rong Lin, Zhengdong D. Zhang

## Abstract

Mosaic loss of chromosome Y (LOY) is a common age-associated somatic alteration in men and is typically measured from DNA-based assays. Many cohorts, however, contain bulk RNA-seq data without matched DNA-based LOY measurements. We developed a Bayesian framework to estimate the fraction of cells with LOY from male bulk RNA-seq by modeling reduced Y-linked gene expression relative to expected expression after adjustment for age, expression covariates, and autosomal/X-linked control genes. In 377 male GTEx samples, individual Y-linked genes showed negative correlations with separately obtained DNA-based LOY measurements, supporting a shared Y-expression depletion signal. The primary fast empirical Bayes estimator achieved a Pearson correlation of 0.678 with measured LOY, a mean absolute error of 1.79%, a root mean squared error of 3.72%, and 95.2% empirical coverage of measured LOY. Performance was strongest for identifying large LOY events, with an AUC of 0.964 for measured LOY greater than 20%, while fine ranking among low-LOY samples remained uncertain. A mixture/PCA hierarchical Bayesian sensitivity model provided similar validation performance and interpretable posterior quantities but did not improve point estimation. Leave-one-Y-gene-out and prior-sensitivity analyses showed that the signal was distributed across multiple Y-linked transcripts and that prior shrinkage affected calibration. In an external whole-blood RNA-seq dataset without measured LOY, estimated LOY showed a modest age-related increase, but ex vivo immune stimulation shifted RNA-derived LOY estimates and reduced multiple Y-linked transcripts, indicating transcriptional confounding. These results show that bulk RNA-seq contains usable information about LOY, especially for larger events, but RNA-derived LOY should be interpreted as a probabilistic transcriptome-based estimate rather than a direct substitute for DNA-based mosaicism measurement.

## INTRODUCTION

Mosaic loss of chromosome Y (mLOY or LOY) is an acquired somatic alteration in which a subset of cells in a male individual lose the Y chromosome. LOY is most readily detected in blood-derived DNA and increases strongly with age, making it one of the most common forms of clonal mosaicism observed in aging men. Large population studies have shown that detectable LOY is frequent in peripheral blood leukocytes, with prevalence rising in older age groups and varying substantially between individuals. Beyond aging itself, epidemiologic and genetic studies have implicated smoking, inherited susceptibility, hematopoietic cell differentiation, and clonal expansion as contributors to LOY development and persistence^1-6^.

Although initially regarded as a largely neutral age-related cytogenetic change, LOY has increasingly been associated with clinically relevant outcomes. Early cohort studies linked LOY in peripheral blood with shorter survival and increased risk of non-hematologic cancer mortality^3^. Subsequent work associated LOY with Alzheimer’s disease and other age-related conditions^1,7^. More recent experimental and human evidence suggests that hematopoietic LOY may contribute directly to disease biology. For example, mouse models and prospective human analyses support a role for hematopoietic Y-chromosome loss in cardiac fibrosis, cardiac dysfunction, and heart failure mortality^8^. These observations have made LOY a useful biomarker of hematopoietic aging and genomic instability, as well as a candidate contributor to male-biased risk for some age-related diseases.

Most LOY studies have relied on DNA-based assays, including SNP-array intensity data^3,4,9,10^, genotype array mosaicism calls^11-14^, and more recent sequencing read-depth patterns or related genomic measurements^15-17^. These approaches are well suited for estimating mosaic chromosomal copy number changes, but they require DNA data of sufficient quality and coverage. In contrast, many human cohorts and public molecular datasets contain bulk RNA-seq but lack matched DNA-based LOY measurements. This creates an opportunity: if Y-chromosome loss reduces the abundance of Y-linked transcripts in bulk tissue, then RNA-seq may contain information about the fraction of cells lacking chromosome Y. However, transcriptomic inference of LOY is not straightforward. Y-gene expression varies by gene, tissue, cell composition, technical factors, and biological state; moreover, inflammatory or immune stimulation can alter Y-gene expression without changing the underlying genomic LOY fraction.

A transcriptome-based LOY estimator thus requires a model that separates Y-chromosome depletion from background expression variation. The key biological signal is conceptually simple: in a bulk sample containing a fraction of LOY cells, expression of Y-linked genes should be reduced relative to the expression expected in cells retaining chromosome Y. In practice, this depletion must be estimated after accounting for age, latent expression structure, technical covariates, and control genes on autosomes and the X chromosome. A useful estimator should also return uncertainty intervals, because low-to-moderate LOY fractions may be difficult to distinguish from ordinary expression variation.

Here, we develop a Bayesian framework to estimate RNA-derived LOY fraction from bulk RNA-seq. The method uses expression of Y-linked genes together with autosomal and X-linked control genes and available covariates to infer a posterior distribution over each individual’s LOY fraction. We implement a fast empirical Bayes estimator for scalable point estimation and a hierarchical Bayesian extension that includes gene-specific Y sensitivity, robust residual errors, mixture priors, age-informed LOY probability, and control-gene factor adjustment. We evaluated the approach in male GTEx samples with measured LOY and further assessed the methods in public blood RNA-seq data without measured LOY, including age-group plausibility and stimulation-induced failure modes.

## MATERIALS AND METHODS

### Study Overview

We developed an RNA-seq method to estimate the fraction of cells with mLOY in male bulk transcriptomes. The approach uses reduced expression of Y-chromosome genes relative to expected expression, after adjustment for covariates and control-gene expression, to infer a latent Y-depletion parameter for each individual. We used separately measured LOY to assess estimation accuracy.

We implemented two main estimators. The primary estimator was a fast empirical Bayes model designed for stable point estimation and scalable application to large cohorts. The secondary estimator was a mixture/PCA hierarchical Bayesian sensitivity model implemented in Stan, designed to quantify uncertainty and evaluate model extensions, including gene-specific Y sensitivity, robust residual errors, mixture priors, age-informed LOY probability, and control-gene factor adjustment.

### Input data and gene sets

The primary validation dataset consisted of bulk RNA-seq expression data from male GTEx whole-blood samples^18^. The input expression matrix contained DESeq2-normalized expression values^19^, donor age, five expression principal components, ten inferred covariates, and a measured LOY fraction (separately measured from DNA sequencing data) used strictly for validation. We used five Y-chromosome genes as the LOY signal genes: DDX3Y, EIF1AY, KDM5D, RPS4Y1, and UTY. We used ten autosomal genes as control genes: IFITM2, FTH1, SRGN, RPLP1, SPI1, NCF2, RPS25, RPL27, FOSL2, and UBL7. We also used five non-pseudoautosomal X-chromosome genes as additional controls: TMSB4X, SAT1, ALAS2, PLP2, and IL2RG. The covariate set included age, the top five expression PCs, the ten inferred covariates, and the autosomal and X-linked control-gene expression values unless otherwise specified. The Y genes were selected a priori as top-expressed non-pseudoautosomal Y-linked transcripts detectable in male bulk RNA-seq. Autosomal and non-PAR X-linked control genes were included as highly expressed comparators and adjustment features, not as invariant housekeeping standards. These controls were not selected by optimizing prediction against measured LOY, and they may themselves be affected by immune activation, aging, or leukocyte composition. For this reason, we evaluated control-gene and covariate ablation analyses and stimulation stress tests.

### Biological measurement model

The model is based on the expectation that, in a bulk sample containing a fraction (*f*_*i*_) of LOY cells, expression of Y-linked genes is reduced relative to the expression expected in cells retaining chromosome Y. We parameterized this depletion on a log2 scale using *λ*_*i*_ = − log_2_(1 − *f*_*i*_) so that 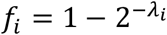. Here, *f*_*i*_ is the estimated cellular fraction of LOY in sample *i*, and *λ*_*i*_ is the latent Y-expression depletion. This parameterization gives a direct biological interpretation: if half of the cells have lost chromosome Y, then *f*_*i*_ = 0.5 and *λ*_*i*_ = 1. For each Y gene *j*, observed expression was modeled as expected baseline expression minus a shared Y-depletion term plus residual noise: *Y*_*ij*_ = baseline_*ij*_ − *λ*_*i*_ + *ε*_*ij*_ . In the hierarchical model, this was generalized to allow gene-specific sensitivity: *Y*_*ij*_ = baseline_*ij*_ − *s*_*j*_*λ*_*i*_ + *ε*_*ij*_, where *s*_*j*_ is the sensitivity of Y gene *j* to cellular LOY fraction.

### Fast empirical Bayes estimator

The fast empirical Bayes estimator proceeds in four steps. First, for each Y gene, we estimated expected baseline expression using ridge regression on covariates and control-gene expression. This produced a predicted non-LOY expression level for each sample and Y gene. The residual for sample *i* and Y gene *j* was defined as observed minus predicted expression. Second, we estimated a reference subset of samples likely to have little or no LOY. The model was initialized using all samples as reference, fit residuals were used to compute a Y-depletion score, and samples with the largest depletion scores were excluded from the reference set. This procedure was iterated to reduce the influence of high-LOY samples on the baseline expression model. Third, we combined residuals across the five Y genes into a single covariance-weighted depletion score. Let **e**_*i*_ be the vector of centered Y-gene residuals for sample *i*, and let ∑ be the shrinkage covariance matrix of residuals estimated in the reference samples. The common Y-depletion score was estimated by projecting the residual vector onto a shared negative Y-expression direction:

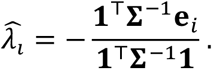

This gives higher weight to Y genes with lower residual variance and accounts for correlation among Y-gene residuals. Fourth, we converted each depletion score into a posterior distribution for *λ*_*i*_. After covariance-weighted projection of the residualized Y-gene vector, the empirical Bayes likelihood was 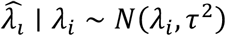, where 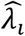 is the generalized least-squares Y-depletion score and *τ* is its standard error estimated from the residual covariance matrix. We used an exponential prior on nonnegative depletion, *λ*_*i*_ ∼ Exp(*r*) with default rate *r* = 20. The posterior was therefore proportional to the product of this Gaussian likelihood and exponential prior, 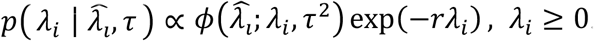, *λ* ≥ 0. Posterior means, medians, and 95% credible intervals were computed by evaluating this posterior on a grid over nonnegative *λ*_*i*_. Finally, *λ*_*i*_ was transformed to the estimated cellular LOY fraction as 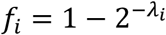. Full details of the covariance-weighted score and grid posterior calculation are provided in Supplementary Methods. The fast empirical Bayes model was used as the primary estimator because it gave the strongest calibrated performance in GTEx validation while remaining computationally simple and stable.

### Hierarchical Bayesian model

We also implemented a hierarchical Bayesian model in Stan^20^. The model jointly estimates sample-specific LOY depletion, gene-specific sensitivity, residual variance, and prior mixture structure. Student-*t* residuals were used to reduce sensitivity to outlying expression values. For sample *i* and Y gene *j*,

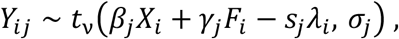

where *X*_*i*_ contains age and other covariates, *F*_*i*_ contains control-gene factor scores, *s*_*j*_ is the sensitivity of Y gene *j*, and *σ*_*j*_ is the gene-specific residual scale. *X*_*i*_ contains the observed and supplied sample-level covariates, including intercept, age, the top five expression PCs, and ten inferred covariates. *F*_*i*_ contains additional control-gene factor scores derived by applying PCA to autosomal and X-linked control-gene expression after residualizing those control genes against *X*_*i*_ . Thus, the PCs in *X*_*i*_ are supplied global expression PCs, whereas the factors in *F*_*i*_ are estimated control-gene residual factors used to capture the remaining control-gene expression structure.

To distinguish near-reference samples from potential LOY samples, the prior on *λ*_*i*_ was specified as a two-component mixture:

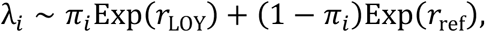

with *π*_*i*_ = logit^−1^(*α*_0_ + *α*Age_*i*_). Here, *r*_ref_ was set larger than *r*_LOY_, so that the reference component was concentrated near zero depletion, while the LOY component allowed larger depletion values. In the default Stan model, we used *r*_ref_ =100 and *r*_LOY_ =10, corresponding to prior mean LOY fractions of approximately 0.7% and 6.5%, respectively. We varied *r*_LOY_ in prior-sensitivity analyses.

The mixture probability *π*_*i*_ was allowed to depend on age, reflecting the known increase of LOY with aging. Control-gene factors were estimated by PCA of residualized autosomal and X-linked control-gene expression and were included to capture latent expression structure not explained by the supplied covariates. The hierarchical model was used as a probabilistic sensitivity model rather than the primary point estimator. It provides interpretable posterior quantities, including gene-specific sensitivity, posterior probability of belonging to the LOY component, and uncertainty intervals for each individual’s LOY fraction.

### Validation strategy

Measured LOY fractions were used only for validation after model fitting. For each method, we compared the estimated LOY fraction with the measured LOY fraction using: Pearson correlation, Spearman correlation, mean absolute error, root mean squared error, bias, 95% credible interval (CI) coverage, AUC for measured LOY above 3%, 5%, 10%, and 20%. All error metrics were reported in percentage points (%) after multiplying LOY fractions by 100. We used bootstrap resampling of individuals to obtain 95% confidence intervals for validation metrics.

### Robustness and sensitivity analyses

We performed several robustness analyses for the fast empirical Bayes estimator. First, we conducted leave-one-Y-gene-out analysis by refitting the model after removing each Y gene in turn. This tested whether performance depended disproportionately on a single Y-linked transcript. Second, we evaluated prior sensitivity by varying the exponential prior rate on *λ*_*i*_: rate = 10, 20, 50, and 100. Third, we performed covariate and control-gene ablation analyses comparing: intercept only, age only, age + PCs + inferred covariates, covariates + autosomal controls, covariates + autosomal + X controls, covariates + control-gene PCA factors. For the hierarchical Stan model, we similarly varied the LOY-component prior rate and assessed accuracy, interval coverage, posterior shrinkage, and sampler diagnostics.

### Application to GSE279480

To evaluate external behavior in a public blood RNA-seq dataset without measured LOY, we applied the method to GSE279480, a whole-blood RNA-seq study of healthy adult immunotypes^21^. This dataset includes two adult age groups, 25-35 and 55-65 years, multiple longitudinal timepoints, and ex vivo stimulation conditions. Raw gene counts from GSE279480_P441_genecounts.csv were mapped to sample metadata using GSE279480_family.soft, where the ‘!Sample_description’ field links GEO sample records to internal library identifiers in the count matrix. Counts were normalized to log_2_(CPM + 1) for the LOY gene panel. Primary LOY estimation was restricted to male unstimulated Null samples. When multiple unstimulated samples were available for the same donor, one sample per donor was selected for the primary analysis, and donor-averaged estimates across repeated Null samples were used as a sensitivity analysis. Because GSE279480 does not include measured LOY, these analyses were treated as biological plausibility and robustness analyses rather than accuracy validation. We compared RNA-estimated LOY between the younger and older male age groups and evaluated within-donor stability across repeated Null samples.

### Stimulation stress test

GSE279480 also includes paired ex vivo stimulation samples using LPS, Poly I:C, and SEB. These stimulations should not change the true genomic fraction of cells with LOY over the assay time scale. Therefore, differences between stimulated samples and matched Null samples were interpreted as evidence of transcriptional confounding in RNA-derived LOY estimates. For each paired donor/timepoint, we compared estimated LOY and Y-gene expression in each stimulation condition against the matched Null sample. This analysis was used to identify whether immune activation can mimic Y-expression depletion and bias transcriptome-based LOY inference.

### Implementation

Analyses were implemented in R. Bayesian hierarchical models were fit using Stan through rstan. Model outputs, validation summaries, robustness tables, and manuscript-support figures were generated using reproducible R scripts. Source code for the primary empirical Bayes estimator and hierarchical Bayesian sensitivity model is available at GitHub (https://github.com/zdz-lab/bulk-rna-seq-loy).

## RESULTS

### RNA-seq contains a Y-chromosome depletion signal associated with LOY

We first evaluated whether bulk RNA-seq contains a measurable Y-chromosome depletion signal associated with independently measured LOY. The primary validation dataset included 377 male GTEx samples with RNA-seq expression data and measured LOY fractions available for validation only. Measured LOY varied substantially across individuals, with a median of 2.10%, a mean of 2.95%, and a maximum of 50.38%. Most samples had low measured LOY, but larger events were present: 83 samples had measured LOY greater than 3%, 31 greater than 5%, 10 greater than 10%, and 5 greater than 20% (**Figure 2A**).

Individual Y-chromosome genes showed the expected negative association with measured LOY. Across the five Y genes, Pearson correlations with measured LOY ranged from −0.148 to −0.230 (**Figure 2B**). The strongest marginal associations were observed for EIF1AY (*r* = −0.230; **Figure 2C**), RPS4Y1 (*r* = −0.219), UTY (*r* = −0.197), and DDX3Y (*r* = −0.197), while KDM5D showed a weaker but still negative association (*r* = −0.148). By comparison, the selected autosomal and non-PAR X-chromosome control genes showed weaker and directionally mixed marginal correlations with measured LOY (**Figure 2B**). The median absolute Pearson correlation was 0.197 for Y genes, compared with 0.098 for autosomal controls and 0.083 for X-linked controls (**Figure 2D**).

These results support the presence of a shared Y-expression depletion signal in bulk RNA-seq. At the same time, the modest gene-level correlations indicate that individual Y genes are noisy proxies for LOY. This motivated a multigene model that estimates a shared Y-depletion parameter after adjustment for covariates and control-gene expression (**Figure 1**; **Table 1**).

**Table 1.** Gene sets and covariates.

| *Group* | *Members* | *Chromosomal category* | *Role in model* | *Used in* |  |
| --- | --- | --- | --- | --- | --- |
|  |  |  |  | *Primary EB* | *Mixture PCA Stan* |
| Y genes | DDX3Y,<br>EIF1AY,<br>KDM5D,<br>RPS4Y1, UTY | Chromosome Y | Signal genes used to infer shared Y-expression depletion | Yes | Yes |
| Autosomal control genes | IFITM2, FTH1,<br>SRGN, RPLP1,<br>SPI1, NCF2,<br>RPS25, RPL27,<br>FOSL2, UBL7 | Autosomal | Control genes for baseline expression and cell-state adjustment | Yes | Indirectly |
| X non-PAR control genes | TMSB4X,<br>SAT1, ALAS2,<br>PLP2, IL2RG | X chromosome, non-PAR | Additional sex-chromosome controls not expected to decrease with LOY | Yes | Indirectly |
| Age | age | Phenotypic covariate | Baseline covariate and age-informed prior input | Yes | Yes |
| Expression PCs | PC1-PC5 | Latent expression covariates | Adjustment for broad expression structure | Yes | Yes |
| Inferred covariates | InfCov1-InfCov10 | Latent technical biological covariates | Adjustment for residual expression/batch structure | Yes | No |
| Control-gene PCA factors | PCA scores from residualized autosomal and X control genes | Derived latent factors | Control-gene latent factor adjustment in mixture/PCA Stan and sensitivity analyses | No | Yes |

**Figure 1.**
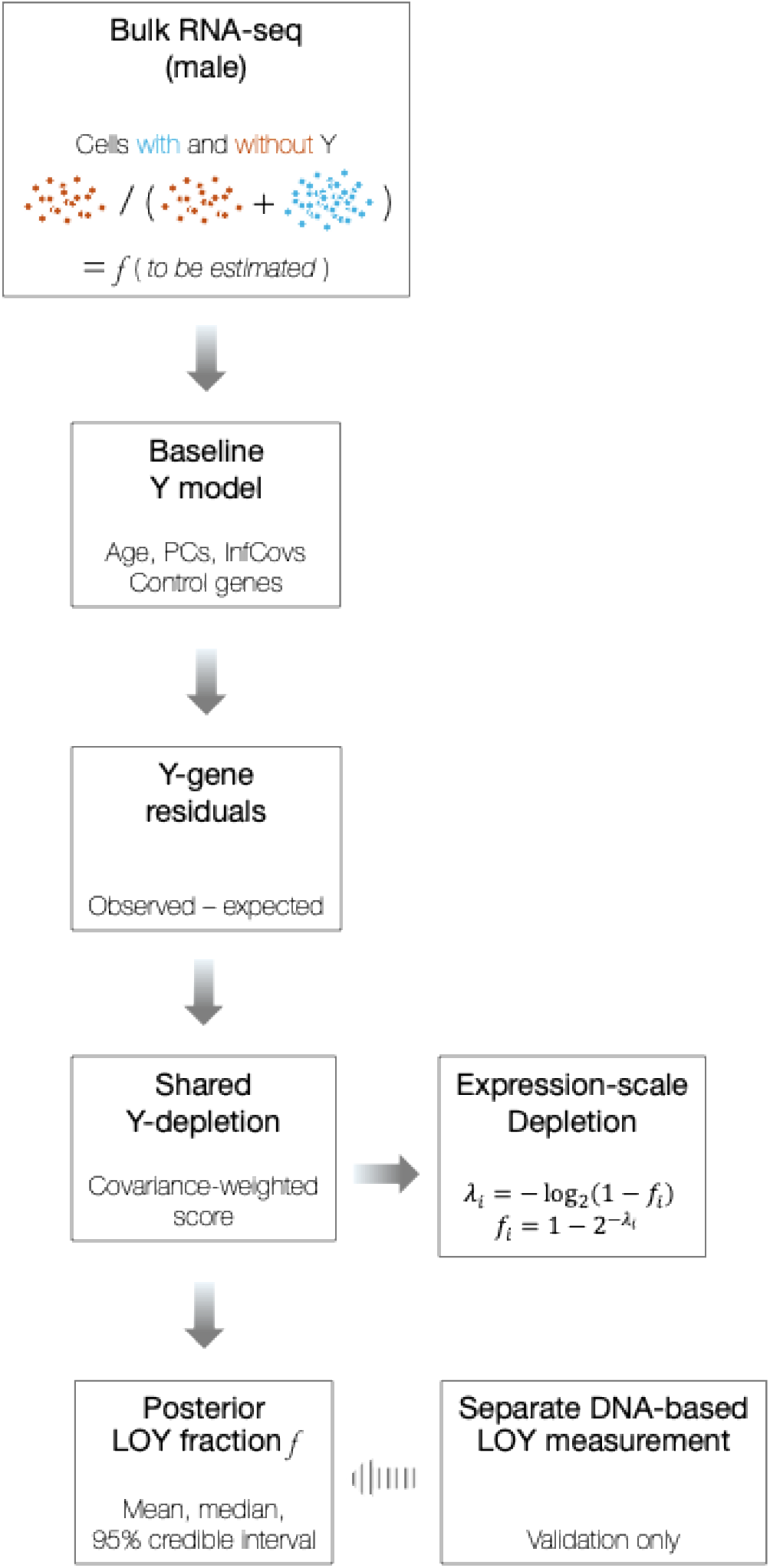
Overview of RNA-seq-based Y-expression depletion estimation. The method models reduced Y-linked gene expression relative to expected expression after covariate and control-gene adjustment. The latent depletion parameter is transformed to an RNA-derived LOY fraction. Measured LOY values are used only for validation.

**Figure 2.**
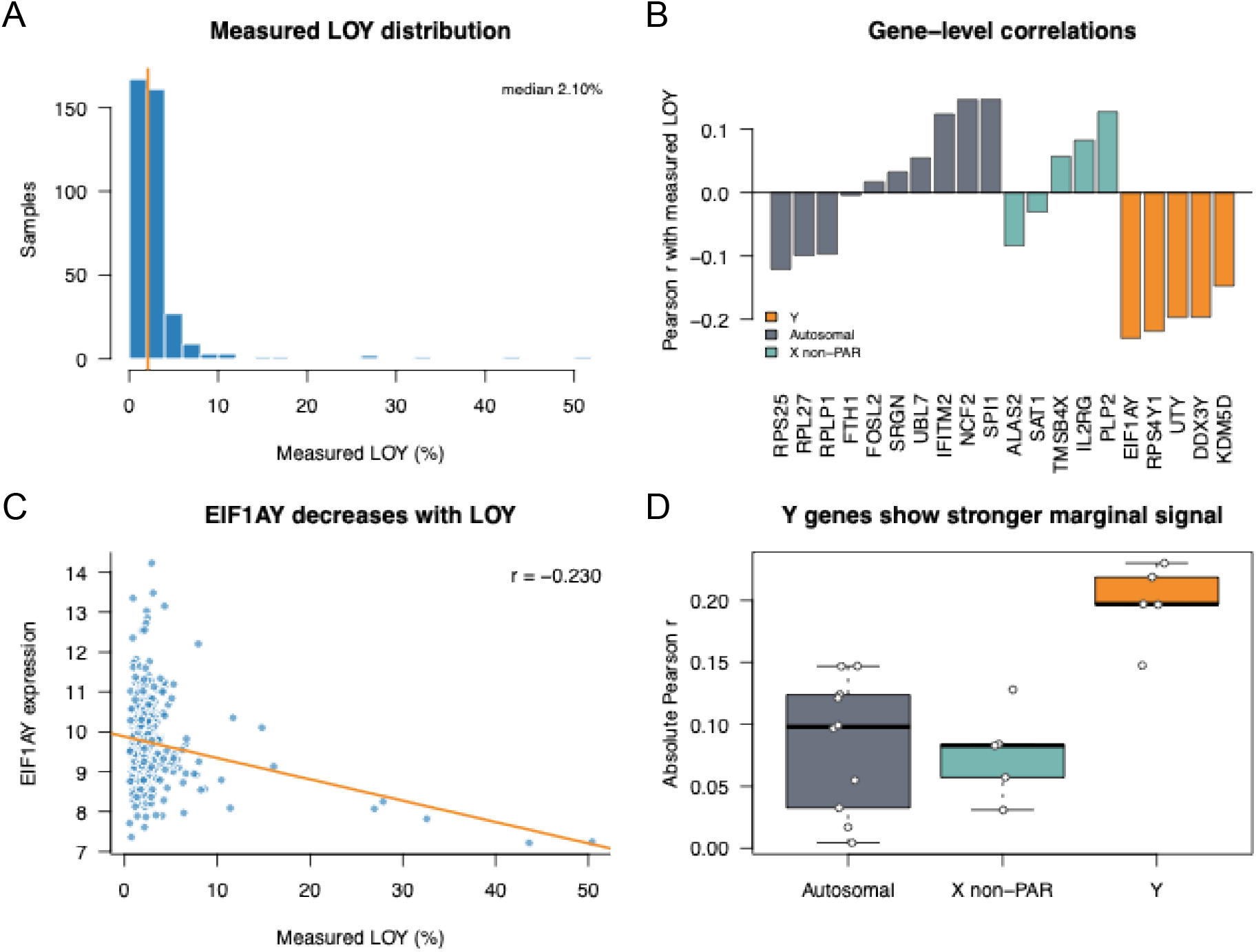
Y-gene expression contains an LOY-associated depletion signal. In 377 male GTEx samples, measured LOY ranged from low-level mosaicism to large events. Individual Y genes showed negative correlations with measured LOY, whereas autosomal and non-PAR X control genes showed weaker and directionally mixed associations. (A) Measured LOY distribution in GTEx males. (B) Correlations between measured LOY and expression of Y genes, autosomal controls, and X controls. (C) Representative Y-gene (EIF1AY) expression vs measured LOY scatterplots. (D) Coefficient plot showing Y genes have the biggest LOY associations.

### Fast empirical Bayes estimation recovers large LOY events and provides calibrated uncertainty

We next evaluated the fast empirical Bayes estimator, which combines residualized Y-gene expression into a covariance-weighted depletion score and converts this score into a posterior distribution for LOY fraction.

In validation against measured LOY, the fast empirical Bayes estimator achieved a Pearson correlation of 0.678 between observed and estimated LOY (**Table 2**; **Figure 3A**; **Supplementary Figure S1**). Bootstrap resampling gave a 95% confidence interval of 0.158 to 0.873. Mean absolute error was 1.79%, and root mean squared error was 3.72%. The estimator showed modest positive bias, with the mean estimated LOY exceeding the measured LOY by 0.42% (**Figure 3B**).

**Table 2.** GTEx validation performance.

| *Method* | *Fast EB* | *Mixture/PCA Stan* |
| --- | --- | --- |
| Uses measured LOY for fitting | No | No |
| Pearson $r$ | 0.678 [0.158, 0.873] | 0.665 [0.203, 0.862] |
| Spearman $\rho$ | 0.004 [−0.095, 0.107] | 0.079 [−0.023, 0.178] |
| MAE (%) | 1.788 [1.486, 2.148] | 1.943 [1.645, 2.298] |
| RMSE (%) | 3.718 [2.456, 5.061] | 3.784 [2.604, 5.005] |
| Bias (%) | 0.418 [0.057, 0.805] | −0.747 [−1.111, −0.383] |
| Coverage 95% | 0.952 [0.931, 0.973] | 0.918 [0.889, 0.944] |
| AUC LOY >10% | 0.726 [0.543, 0.901] | 0.731 [0.549, 0.890] |
| AUC LOY >20% | 0.964 [0.883, 1.000] | 0.948 [0.833, 1.000] |

Note: Bootstrap 95% credible intervals (CIs) are shown in square brackets.

**Figure 3.**
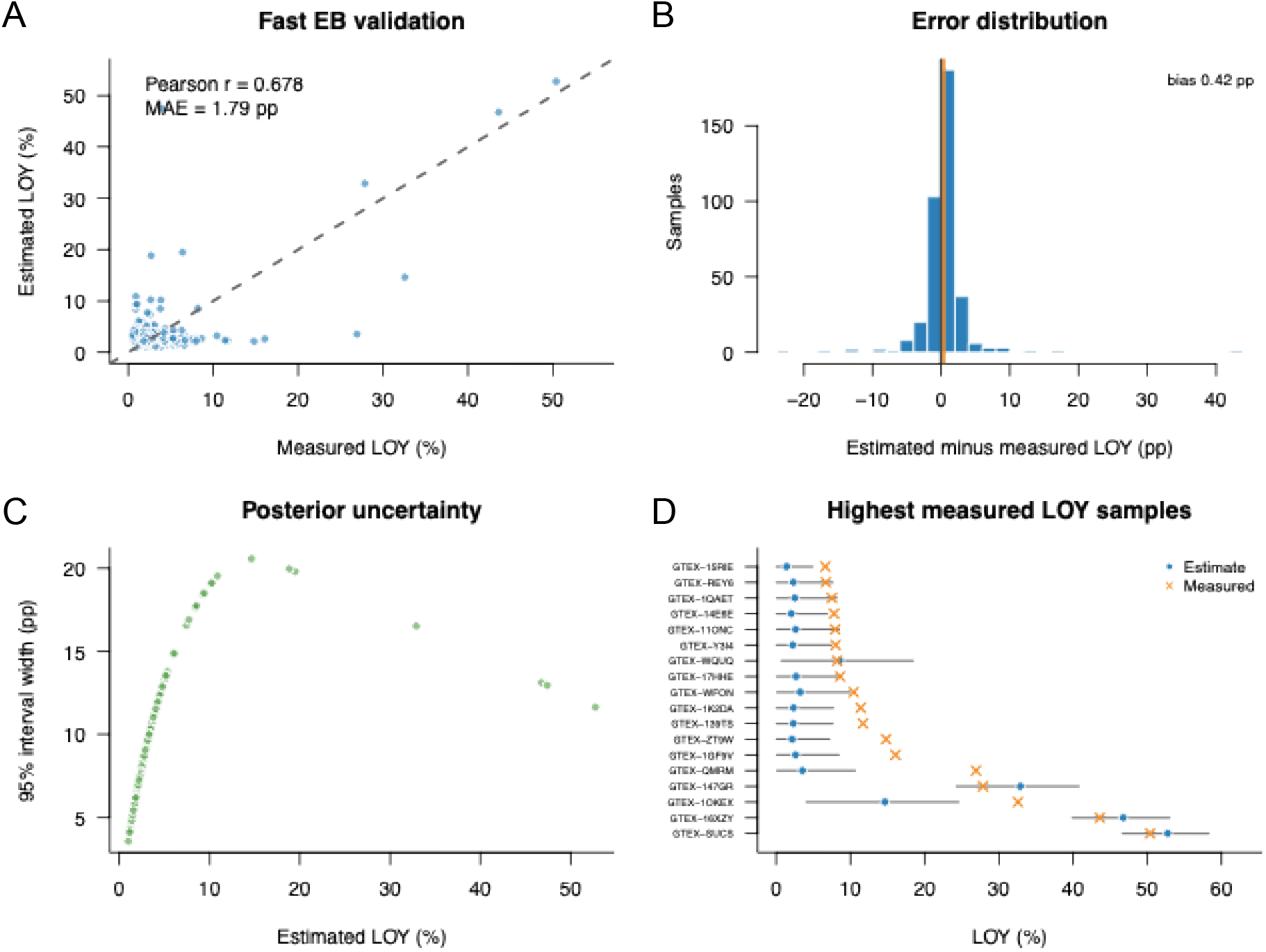
Fast empirical Bayes estimator validation. The empirical Bayes estimator was fit without using measured LOY. In validation, estimated LOY correlated with measured LOY and showed strong discrimination of large LOY events. Posterior intervals were broad, reflecting limited transcriptomic information at low LOY fractions, but achieved near-nominal empirical coverage. (A) Observed vs estimated LOY scatterplot for fast EB. (B) Error distribution in percentage points. (C) Posterior interval width vs estimated LOY. (D) Calibration/coverage summary or observed LOY with 95% credible intervals for selected samples.

The estimator performed particularly well for identifying large LOY events. The AUC for measured LOY greater than 20% was 0.964, with a bootstrap 95% confidence interval of 0.883 to 1.000 (**Table 2**; **Supplementary Table S1**). In contrast, rank-order performance across the full low-to-high LOY range was limited, with Spearman correlation close to zero (**Table 2**). Thus, the RNA-seq signal was most informative for identifying larger Y-loss events, whereas fine ranking among low-LOY samples remained uncertain. Additional low-LOY and high-outlier diagnostics clarified the source of this performance. When analysis was restricted to samples with measured LOY ≤10%, the fast empirical Bayes estimator had Pearson *r* = 0.096 and Spearman *ρ* = −0.034; when restricted to measured LOY ≤5%, Pearson *r* = 0.084 and Spearman *ρ* = −0.045. Excluding the five samples with measured LOY >20% reduced the Pearson correlation from 0.678 to 0.057. Thus, the overall Pearson correlation was driven largely by the small number of high-LOY samples, and the estimator should not be interpreted as providing a reliable monotonic ranking among individuals with low measured LOY (**Supplementary Figure S5**; **Supplementary Table S5**).

The empirical Bayes model also produced posterior credible intervals for each individual. The median 95% interval width was 8.10%, reflecting the limited information content of a small number of Y-linked transcripts (**Figure 3C**). Empirical coverage of the measured LOY fraction was 95.2%, close to nominal (**Table 2**). This supported the empirical Bayes estimator as a calibrated primary estimator: it provided competitive point-estimation performance, strong detection of large LOY events, and credible intervals that appropriately reflected uncertainty (**Table 2**; **Figure 3D**).

To determine whether the estimator depended disproportionately on one Y gene, we repeated the analysis after removing each Y gene in turn. Performance was stable across all leave-one-gene-out analyses (**Figure 4B**; **Supplementary Figure S3**). Pearson correlation remained between 0.654 and 0.676 after dropping any single Y gene, compared with 0.678 using all five Y genes. RMSE ranged from 3.53 to 3.99% (**Table 3**; **Supplementary Figure S3**). Dropping RPS4Y1 had the largest effect on high-LOY classification, reducing AUC for measured LOY greater than 20% from 0.964 to 0.875 (**Figure 4B**; **Supplementary Figure S3**). Nevertheless, the overall estimator remained functional, indicating that the LOY signal is distributed across multiple Y-linked transcripts rather than being driven by a single gene.

**Table 3.**
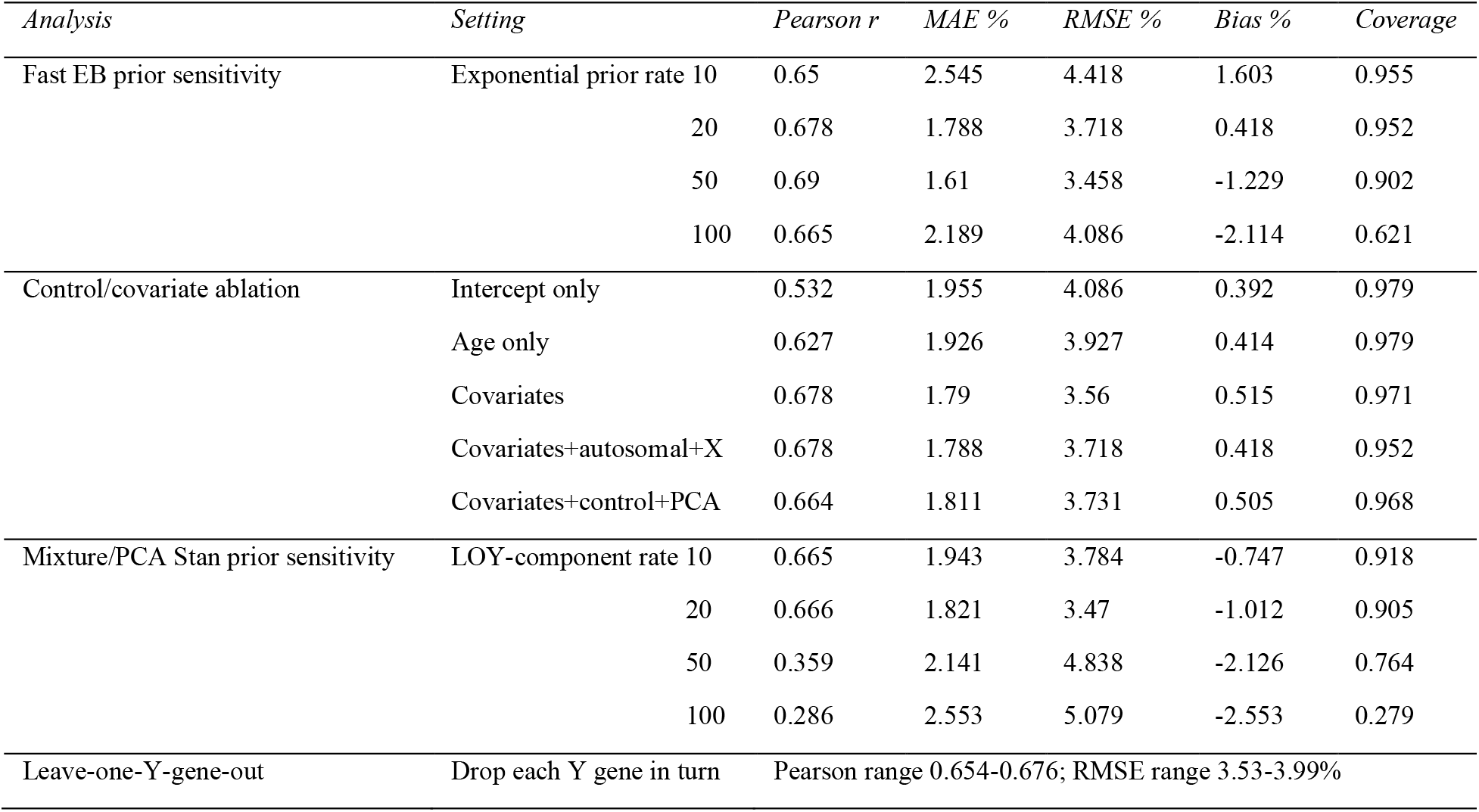
Robustness and sensitivity.

**Figure 4.**
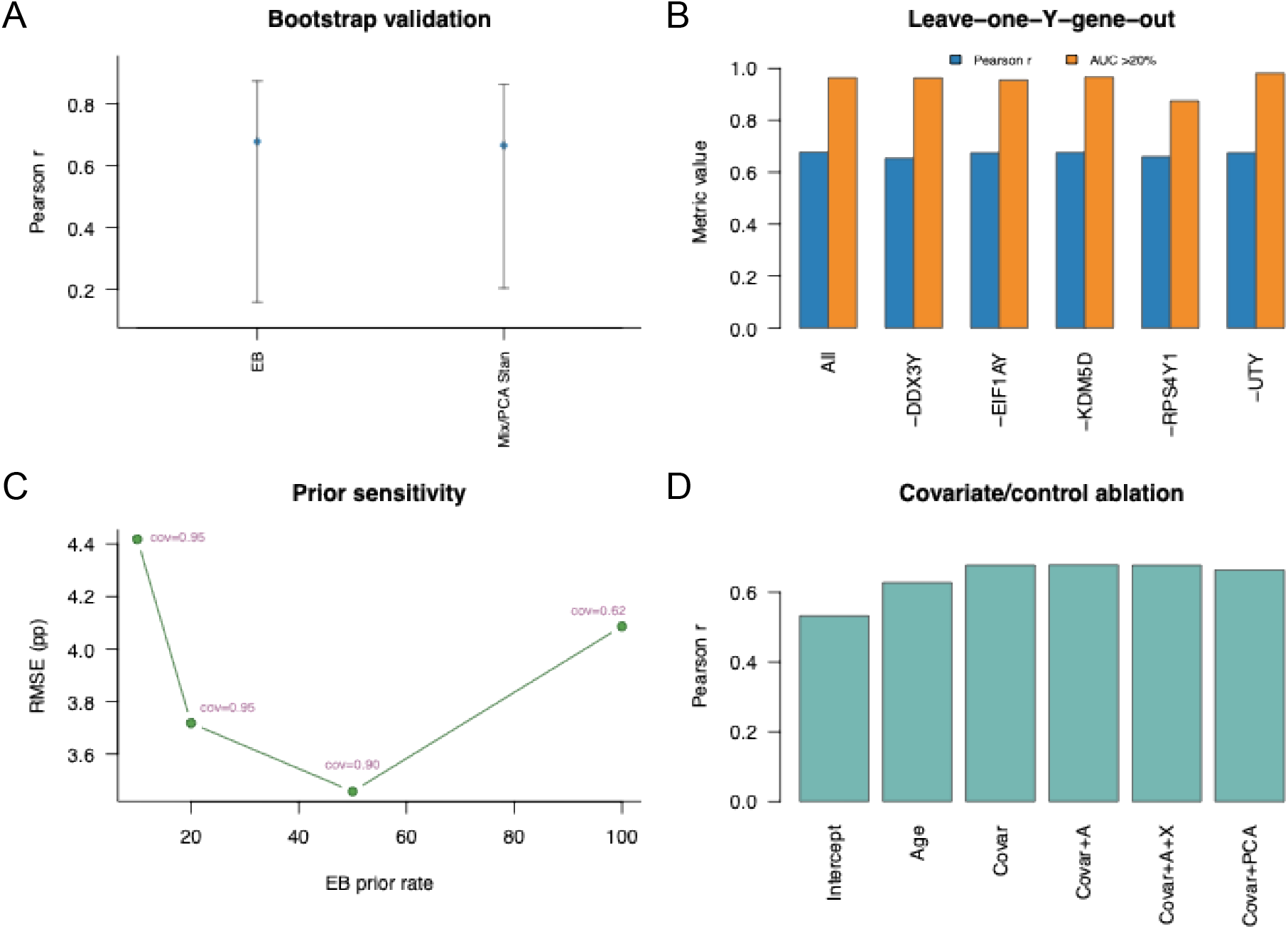
Model comparison and robustness analyses. Bootstrap validation showed that the fast empirical Bayes estimator provided the best calibrated primary method, while the mixture/PCA Stan model served as a conservative Bayesian sensitivity model. Leave-one-Y-gene-out analyses showed that performance was not driven by a single Y gene. Prior sensitivity analyses showed that stronger shrinkage can improve point-error metrics but reduces coverage and underestimates true LOY. (A) Bootstrap comparison of Pearson *r*, MAE, RMSE, and AUC >20% LOY for EB and Stan models. (B) Leave-one-Y-gene-out performance. (C) EB prior sensitivity, showing point-error and coverage tradeoff. (D) Covariate/control-gene ablation.

### Bayesian sensitivity model and prior analyses define the calibration tradeoff

We evaluated the mixture/PCA-factor Stan model, a hierarchical Bayesian extension, to determine whether a more complete probabilistic model improved inference or provided useful sensitivity analyses. This Bayesian model achieved a Pearson correlation of 0.665, MAE of 1.94%, RMSE of 3.78%, and a coverage of 91.8% (**Table 2**; **Figure 4A**; **Supplementary Figure S1**). It was more conservative than the empirical Bayes estimator, with a negative bias of −0.75%, but it retained strong discrimination of large LOY events, with an AUC of 0.948 for measured LOY greater than 20% (**Table 2**). Although the mixture/PCA Stan model did not outperform the empirical Bayes estimator as a primary point estimator, it provided interpretable posterior quantities and useful sensitivity checks.

The mixture/PCA model estimated gene-specific Y sensitivity, sample-level LOY-component probability, and control-gene factor effects, thereby providing a richer probabilistic representation of the RNA-derived LOY signal (**Supplementary Table S2**). For example, gene-specific LOY sensitivities were positive for all five Y genes, supporting a shared Y-depletion direction. Posterior mean sensitivities were highest for UTY (1.59; 95% CI, 1.26-1.97), EIF1AY (1.51; 95% CI, 0.96-2.14), and DDX3Y (1.42; 95% CI, 0.99-1.88), and lower for KDM5D (1.05; 95% CI, 0.75-1.40) and RPS4Y1 (1.03; 95% CI, 0.80-1.32). The age coefficient in the mixture prior was positive (*α* = 0.87; 95% CI, 0.17-1.68), indicating that older samples had a higher prior probability of belonging to the LOY component. The implied LOY-component probability increased from approximately 0.10 at one standard deviation below the mean age to 0.34 at one standard deviation above the mean age. These posterior summaries support the model structure: LOY is captured primarily by a shared positive depletion factor, while age contributes both to LOY-component probability and to residual gene-specific baseline expression.

We retained the empirical Bayes estimator as the primary method and used the mixture/PCA Stan model as the Bayesian sensitivity model (**Table 2**). Prior sensitivity analyses showed that prior strength strongly affected calibration. For the empirical Bayes estimator, a weaker exponential prior on the depletion parameter, rate 10, increased positive bias and worsened point-estimation error, with MAE 2.55% and RMSE 4.42% (**Table 3**; **Figure 4C**). The default rate of 20 gave a favorable balance of error and calibration, with MAE 1.79%, RMSE 3.72%, bias 0.42%, and coverage 95.2%. A stronger prior, rate 50, improved point-error metrics, with MAE 1.61% and RMSE 3.46%, but introduced negative bias (−1.23%) and reduced coverage to 90.2%. An even stronger prior, rate 100, over-shrank estimates toward zero, producing bias of -2.11% and coverage of only 62.1%.

Range-stratified calibration showed the same pattern. In samples with measured LOY ≤3%, the mean estimated LOY was 2.84% compared with the mean measured LOY of 1.82%, whereas estimates were lower than measured LOY in the higher but still sparse LOY strata. In the five samples with measured LOY >20%, the mean estimated LOY was 30.11% compared with the mean measured LOY of 36.27% (**Supplementary Table S6**). These results support the use of the method primarily as an uncertainty-aware RNA-derived screen for larger Y-depletion events rather than as a precise continuous estimator across the low-LOY range.

The mixture/PCA Stan model showed a similar sensitivity to overly strong shrinkage. LOY-component prior rates of 10 and 20 produced comparable Pearson correlations (0.665 and 0.666, respectively), while rate 20 improved RMSE relative to rate 10 (**Supplementary Figure S2**). However, rates 50 and 100 substantially over-shrank estimates, reducing Pearson correlation to 0.359 and 0.286, respectively, and lowering interval coverage to 76.4% and 27.9%. All Stan prior-sensitivity fits had zero divergent transitions, with maximum 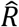 values near 1. These results indicate that stronger priors can improve apparent point-estimation error in low-LOY-dominated samples but at the cost of underestimating true LOY and losing interval calibration. We therefore used the empirical Bayes rate 20 as the calibrated default and treated stronger shrinkage settings as sensitivity analyses.

We also assessed the contribution of covariates and control genes. An intercept-only empirical Bayes model achieved a Pearson correlation of 0.532 and RMSE of 4.09% (**Table 3**; **Figure 4D**; **Supplementary Figure S4**). Including age alone improved Pearson correlation to 0.627. Including age, expression PCs, and inferred covariates increased Pearson correlation to 0.678 and reduced RMSE to 3.56%. Adding autosomal and X-linked control genes did not materially improve Pearson correlation in GTEx, but performance remained similar to the covariate-only model. The full covariate plus autosomal plus X-control specification achieved a Pearson correlation of 0.678 and RMSE of 3.72%. A control-gene PCA version achieved a Pearson correlation of 0.664 and RMSE of 3.73%.

These analyses show that age and latent expression covariates are important for modeling baseline Y-gene expression. Control-gene adjustment did not substantially improve GTEx validation metrics, but it provides a biologically motivated adjustment layer that may be useful for external datasets where batch effects, cell composition, or cell-state structure differ from the validation cohort.

### Blood RNA-seq application reveals biological plausibility and stimulation-related confounding

Finally, we applied the method to GSE279480, a public whole-blood RNA-seq dataset from healthy adults without measured LOY. Because no DNA-based LOY measurements were available, this analysis was treated as an external biological plausibility and robustness assessment rather than an accuracy validation. We restricted the primary analysis to male unstimulated Null samples and selected one sample per donor, yielding 43 male donors: 21 aged 25-35 years and 22 aged 55-65 years (**Figure 5**; **Table 4**; **Supplementary Tables S3a-b**).

**Table 4.** LOY estimation of GSE279480.

| *Analysis* | *Comparison* | *N* | *Estimator* | *Difference (%)* |  | *P value* |
| --- | --- | --- | --- | --- | --- | --- |
|  |  |  |  | *Mean* | *Median* |  |
| Age group comparison | 55-65 vs 25-35 | 22 older, 21 younger | Fast EB | 0.379 | 0.312 | 0.355 |
|  |  |  | Mixture/PCA Stan | 0.468 | 0.411 | 0.00402 |
| Stimulation stress test | LPS vs Null | 109 | Fast EB | 0.293 | 0.168 | 0.0162 |
|  | Poly I:C vs Null | 109 | Fast EB | 1.227 | 0.428 | 3.32E-06 |
|  | SEB vs Null | 110 | Fast EB | 0.706 | 0.295 | 0.000645 |

**Figure 5.**
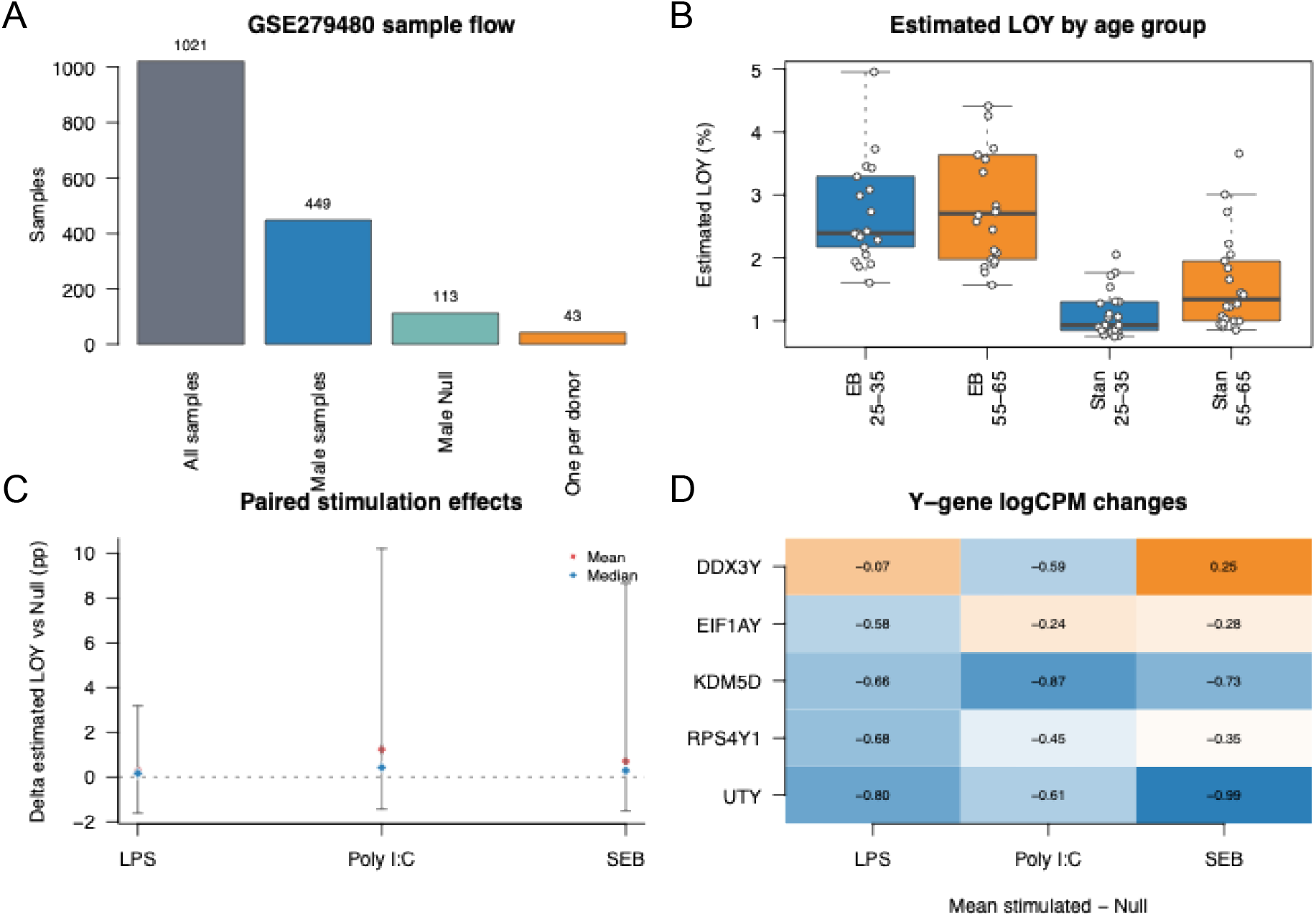
Application to GSE279480 and stimulation stress test. In GSE279480, male unstimulated samples showed a modest age-related increase in RNA-estimated LOY, especially with the mixture/PCA Stan model. Because measured LOY was unavailable, this analysis was interpreted as biological plausibility rather than validation. Paired stimulation analyses showed that LPS, Poly I:C, and SEB shifted RNA-derived LOY estimates relative to matched Null samples, indicating that immune activation can mimic Y-expression depletion. (A) GSE279480 sample flow: total samples, male samples, Null samples, one-per-donor primary set. (B) Estimated LOY by age group, 25-35 vs 55-65, using male Null samples. (C) Paired stimulation effects on estimated LOY: LPS, Poly I:C, SEB vs matched Null. (D) Paired Y-gene logCPM changes under stimulation, highlighting Poly I:C.

Using the empirical Bayes estimator, the older group showed a small, directionally consistent increase in estimated LOY. Mean estimated LOY was 3.00% in the 25-35 group and 3.38% in the 55-65 group, a difference of 0.38%; this difference was not statistically significant by one-sided Wilcoxon test (*P* = 0.355) (**Figure 5A**; **Table 4**). Using the mixture/PCA Stan model, the older group showed a clearer increase. Mean estimated LOY was 1.12% in the 25-35 group and 1.59% in the 55-65 group, a difference of 0.47% (*P* = 0.004) (**Figure 5B**; **Table 4**). When repeated Null samples were averaged per donor, the same qualitative pattern was observed: the Stan model showed a modest older-group increase, whereas the empirical Bayes model showed only a weak directional difference (**Table 4**). These results are consistent with the known age association of LOY, but because measured LOY was unavailable, they should be interpreted as plausibility evidence rather than validation.

GSE279480 also provided paired ex vivo stimulation samples, allowing us to test whether immune activation alters RNA-derived LOY estimates. Since short-term stimulation should not change the true genomic fraction of LOY cells, differences between stimulated samples and paired Null samples reflect transcriptional confounding rather than true LOY change. Compared with matched Null samples, LPS stimulation increased the empirical Bayes estimated LOY by a mean of 0.29% and a median of 0.17% (Wilcoxon *P* = 0.016) (**Figure 5C**; **Table 4**). Poly I:C produced a larger shift, increasing estimated LOY by a mean of 1.23% and a median of 0.43% (*P* = 3.3E−6). SEB also increased estimated LOY, with a mean shift of 0.71% and a median shift of 0.29% (*P* = 6.5E−4).

At the gene-expression level, stimulation altered multiple Y-linked transcripts. Poly I:C reduced all five Y-gene logCPM values relative to paired Null samples, directly mimicking a Y-expression depletion pattern (**Figure 5D**; **Supplementary Table S4**). These findings identify immune activation as an important potential confounder of RNA-derived LOY inference. They also support restricting primary analyses to baseline or unstimulated samples unless condition effects are explicitly modeled.

Overall, these results show that bulk RNA-seq contains a detectable Y-expression depletion signal that can be converted into an uncertainty-aware estimate of LOY fraction without training on measured LOY. The method is strongest for identifying larger LOY events, less reliable for fine ranking among low-LOY samples, and sensitive to biological perturbations that alter Y-gene transcription.

## DISCUSSION

This study shows that bulk RNA-seq contains a measurable Y-expression depletion phenotype associated with independently measured mosaic loss of chromosome Y. The method should therefore be interpreted as an RNA-derived Y-depletion estimator calibrated to LOY under the biological and technical conditions represented in the validation cohort, rather than as a direct molecular assay of chromosome Y copy number. Reduced expression of Y-linked genes can arise from true genomic LOY, but it can also arise from immune activation, cell-state transitions, leukocyte composition, differentiation, stress responses, RNA quality differences, or sex-biased transcriptional regulation. This distinction is central to the interpretation of the method.

The empirical Bayes estimator performed best as the primary point estimator. Its advantage likely reflects a favorable bias-variance tradeoff for this problem: only a small number of Y-linked transcripts are informative, and their expression is affected by cell composition, technical variation, and biological state. The model therefore benefits from shrinkage, covariance weighting, and residualization against measured and inferred expression covariates. The mixture/PCA Stan model was valuable as a Bayesian sensitivity analysis because it made the prior structure, gene-specific sensitivity, latent control-gene factors, and interval calibration explicit, but the added flexibility did not improve point-estimation accuracy in the available validation data. This result argues for presenting the fast empirical Bayes estimator as the operational method and the mixture/PCA hierarchical Bayesian model as a confirmatory model for uncertainty and prior sensitivity.

A major strength of the framework is that measured LOY was reserved strictly for validation. This makes the approach suitable for cohorts in which matched DNA-based LOY calls are unavailable, including legacy RNA-seq collections and studies where biospecimen constraints make additional DNA profiling impractical. At the same time, the method depends on transcript abundance rather than chromosome copy number. For that reason, it is inherently more vulnerable than DNA-based assays to transcriptional regulation, sample composition, RNA quality, normalization choices, and context-specific regulation of Y-linked genes. The use of DESeq2-normalized GTEx expression data^10,11^ and latent expression adjustment^22^ helps address some of these sources of variation, but it cannot eliminate them.

The external analysis of GSE279480 illustrates both the promise and the caution required for RNA-derived LOY inference. In unstimulated whole-blood samples, the older age group showed a modest increase in estimated LOY, consistent with the well-established age dependence of LOY^3,4,6,7^. However, ex vivo stimulation shifted the estimated LOY even though short-term stimulation should not change the underlying genomic LOY fraction. Poly I:C, in particular, reduced multiple Y-linked transcripts and produced an apparent increase in RNA-derived LOY. These results suggest that inflammatory activation, antiviral response, or other strong transcriptional perturbations can mimic a Y-depletion signature. Applications to disease cohorts should therefore prioritize baseline samples, include disease or stimulation covariates when available, and interpret case-control differences cautiously when immune activation differs across groups.

The paired stimulation analysis provides a useful negative-control stress test. Poly I:C, LPS, and SEB stimulation shifted RNA-derived LOY estimates over a short time scale in which genomic LOY fractions should not plausibly change. These shifts demonstrate that the estimator can detect transcriptional Y-depletion states that mimic LOY. Consequently, estimates from stimulated, inflamed, or compositionally shifted samples should be interpreted as apparent RNA-derived Y depletion unless supported by DNA-based LOY measurements or explicit modeling of condition and cell composition.

These findings complement, rather than replace, the large DNA-based literature on LOY and clonal mosaicism. DNA studies have established that LOY increases with age, is influenced by inherited and environmental factors, and is associated with cancer, mortality, Alzheimer’s disease, cardiovascular outcomes, and broader clonal hematopoietic processes^1-8,13,16^. RNA-seq adds a different layer: it can expose downstream transcriptional consequences and can be applied to cohorts where RNA is the only available molecular assay. The tradeoff is that RNA-derived estimates conflate chromosome dosage with transcriptional state unless the model and study design explicitly account for context. Thus, the most natural use case is not definitive LOY calling in isolation, but hypothesis generation, sample stratification, and sensitivity analysis in transcriptomic cohorts.

Several limitations remain. First, the validation dataset contains relatively few large LOY events, so estimates of performance at high LOY have wide uncertainty. Second, the Y-gene panel is small, and each gene has its own regulation and measurement noise; leave-one-gene-out analysis showed robustness, but also indicated that some discrimination depends on particular transcripts. Third, the current model does not explicitly deconvolve leukocyte subsets, even though LOY prevalence and Y-gene expression may vary across cell types. Fourth, external validation in GSE279480 lacked DNA-based LOY measurements, so its age and stimulation analyses assess biological plausibility and confounding rather than accuracy.

Cell composition is likely a major source of confounding in bulk blood RNA-seq. Different leukocyte subsets may differ in Y-gene expression, true LOY prevalence, and response to stimulation, and bulk expression averages these effects. The present implementation adjusts for age, expression PCs, inferred covariates, and control-gene factors, but it does not explicitly deconvolve cell types. Future versions should incorporate cell-type proportion estimates, subset-specific Y-expression baselines, or paired DNA/RNA data to distinguish genomic LOY from transcriptional and compositional Y-depletion. Future work should evaluate the method in cohorts with matched DNA and RNA from the same blood draw, extend the model to cell-composition adjustment, and test whether transcriptome-derived LOY improves analysis of aging and disease phenotypes beyond standard RNA-seq covariates.

In summary, bulk RNA-seq carries a measurable LOY-related Y-expression depletion signal. A fast empirical Bayes model can convert that signal into calibrated individual-level estimates while preserving measured LOY for validation only. The method is most useful for detecting larger LOY events and for extending LOY analyses to RNA-seq cohorts lacking matched DNA, provided that immune activation and other transcriptional perturbations are treated as potential confounders.

## ACKNOWLEDGMENTS

This research was funded by U19 AG056278, R56 AG088624, R21 AG093047 and R01 AG044829 from the National Institute on Aging, National Institutes of Health. The content is solely the responsibility of the authors and does not necessarily represent the official views of the National Institutes of Health.

## SUPPLEMENTARY MATERIALS

## SUPPLEMENTARY METHODS

### Fast empirical Bayes estimator

For sample *i*, let **e**_*i*_ denote the vector of residualized Y-gene expression values after covariate adjustment and centering relative to the reference samples. Under the empirical Bayes measurement model,

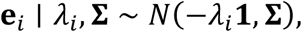

where *λ*_*i*_ ≥ 0 is the latent Y-expression depletion on the log2 scale, **1** is a vector of ones across the Y genes, and **∑** is the residual covariance matrix estimated from the reference samples.

The generalized least-squares estimate of *λ*_*i*_ is

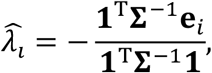

with standard error:

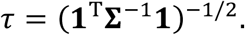

Equivalently, after projecting the multigene residual vector into the one-dimensional depletion score, the likelihood is

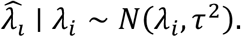

We used an exponential prior on nonnegative depletion,

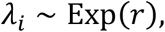

with default rate *r* = 20.

Therefore, the posterior density was

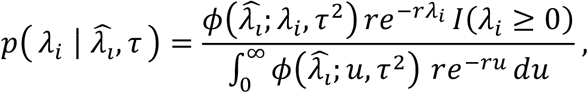

where *ϕ*(⋅; *μ, σ*^2^) denotes the normal density.

In implementation, this posterior was evaluated on a uniform grid of *λ*_*i*_ values from 0 to 2. For grid point *λ*_*k*_, the normalized posterior weight was

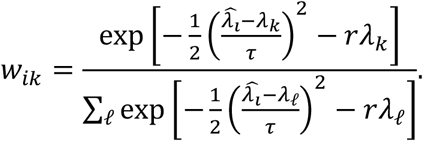

Posterior means, medians, and 95% credible intervals were computed from these grid weights. The depletion parameter was transformed to the cellular LOY fraction by

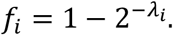

## SUPPLEMENTARY FIGURES

**Supplementary Figure S1.**
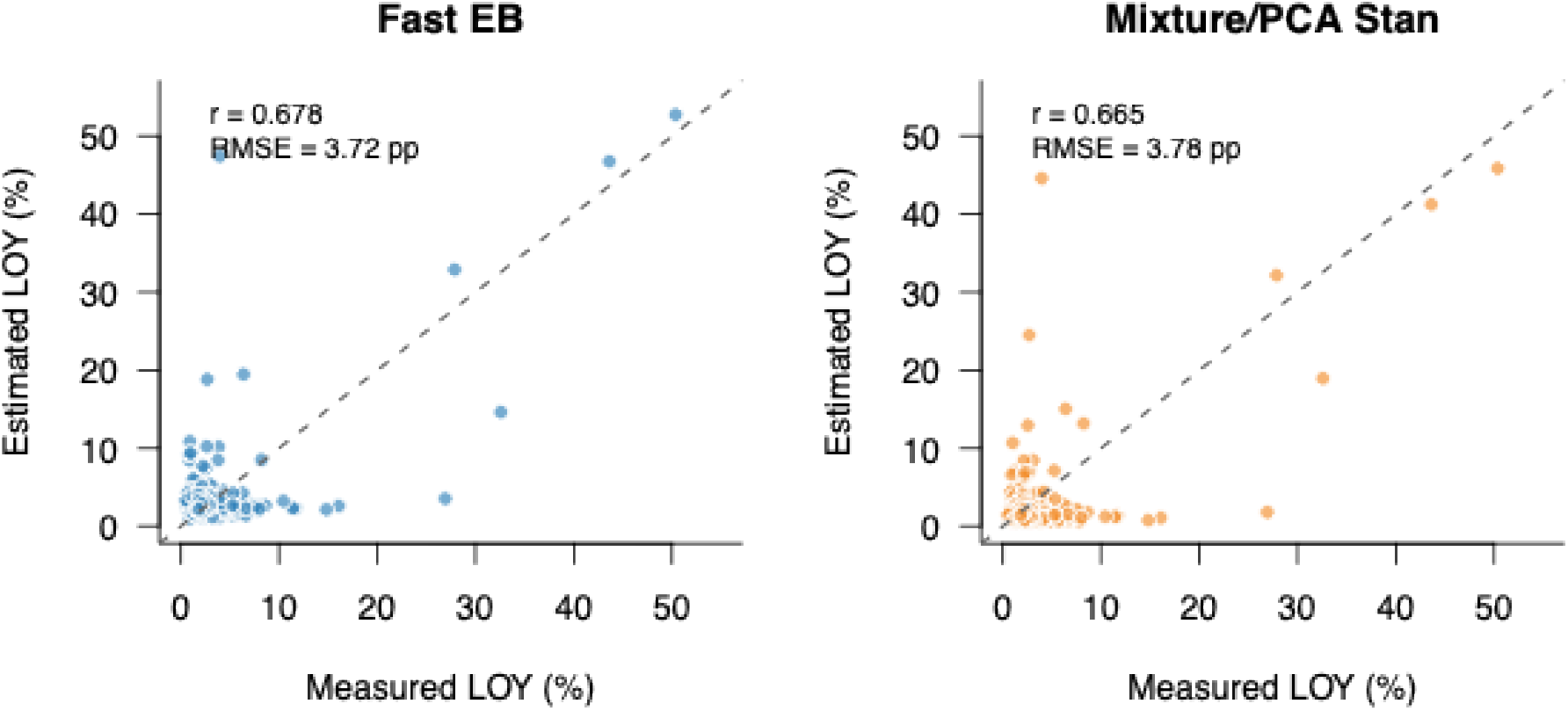
Observed vs estimated LOY for main model variants. Scatterplots compare measured LOY with RNA-estimated LOY for the fast empirical Bayes estimator and mixture/PCA Stan model. Measured LOY was used only for validation.

**Supplementary Figure S2.**
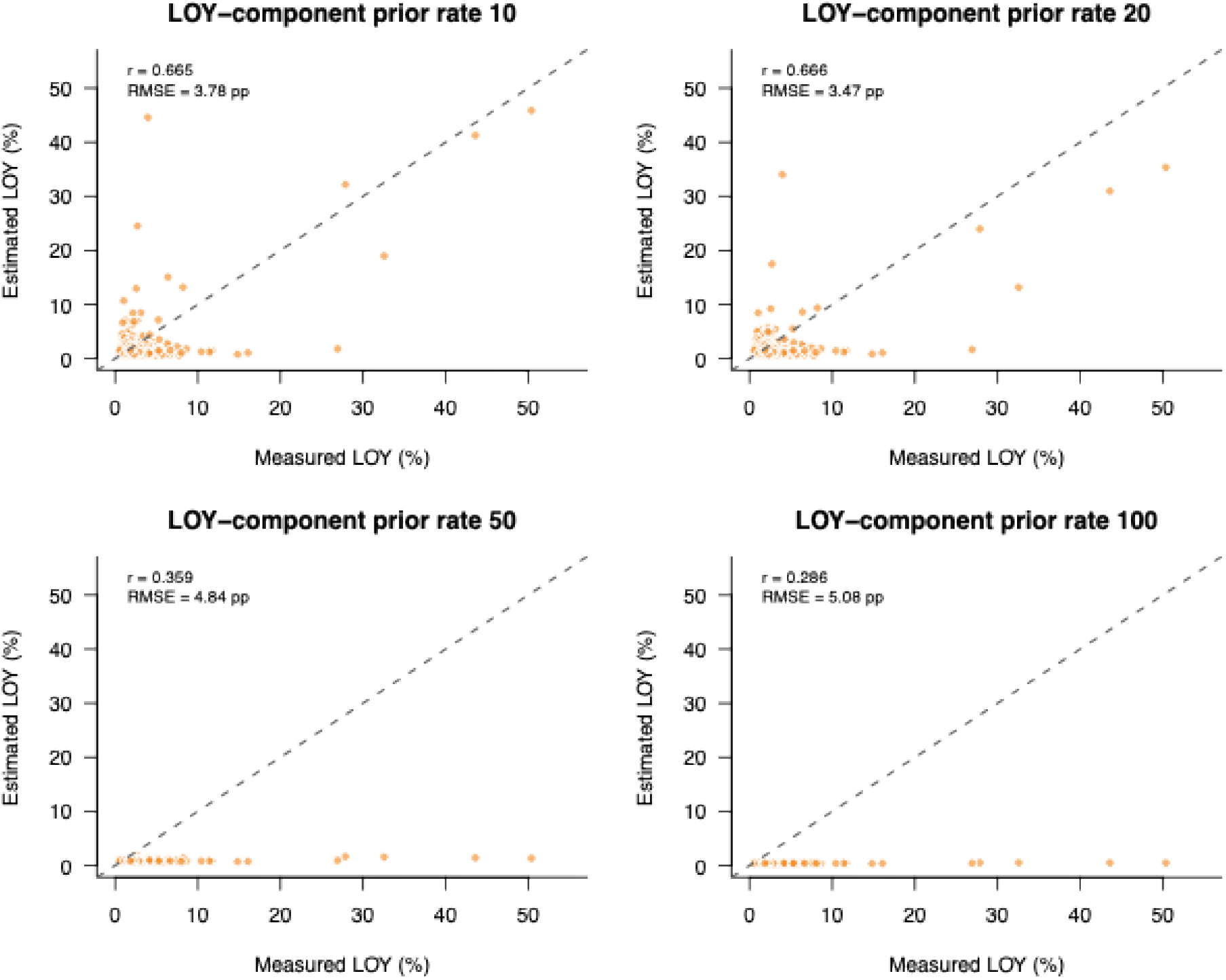
Mixture/PCA Stan prior-sensitivity scatterplots. Observed vs estimated LOY scatterplots for mixture/PCA Stan models fit under LOY-component prior rates 10, 20, 50, and 100. Stronger priors shrink estimates toward zero and reduce recovery of high-LOY samples.

**Supplementary Figure S3.**
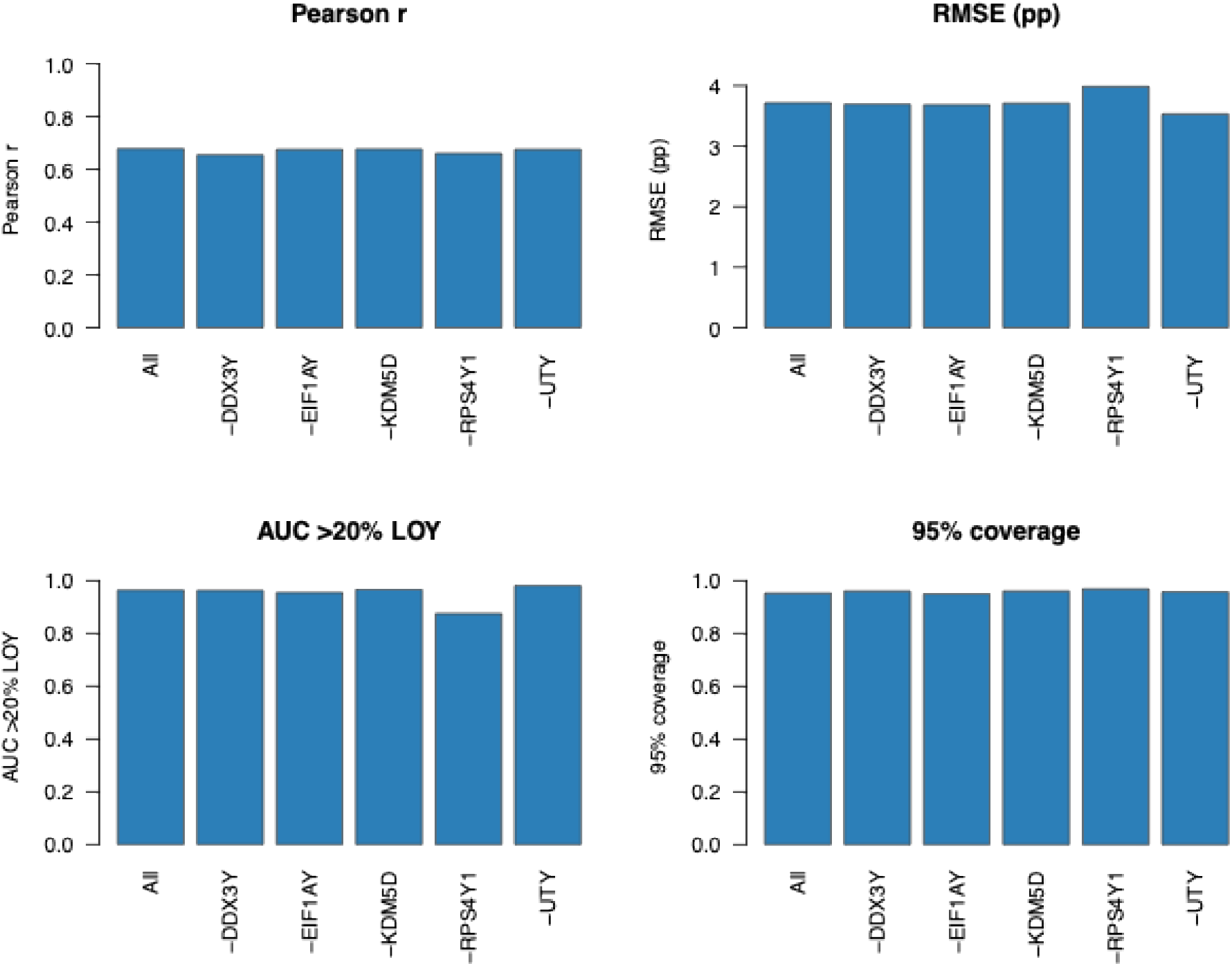
Leave-one-Y-gene-out detailed metrics. Fast empirical Bayes validation metrics after dropping each Y gene in turn. Performance remains stable across leave-one-gene-out analyses, indicating that the signal is not driven by a single Y transcript.

**Supplementary Figure S4.**
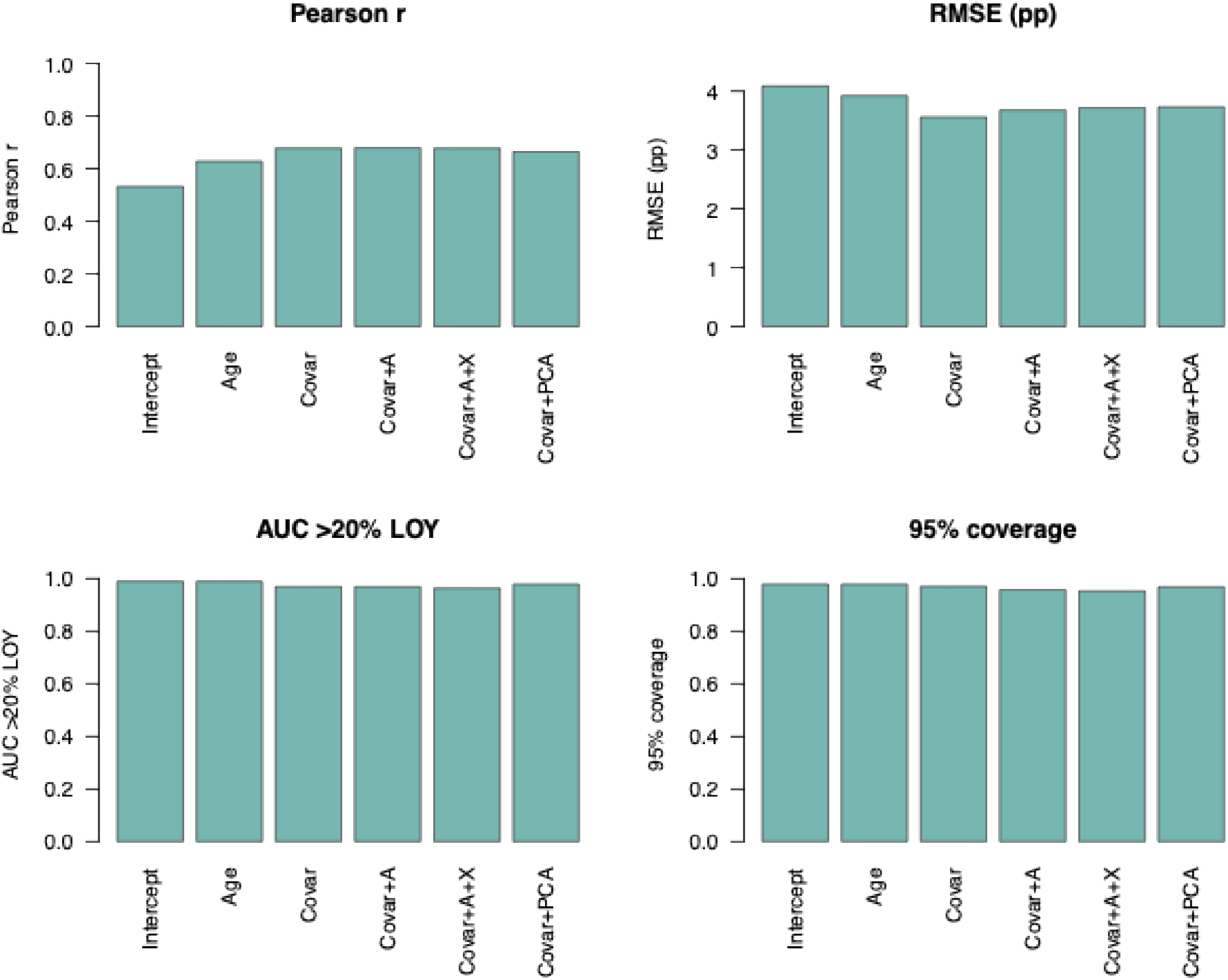
Control-gene/covariate ablation detailed metrics. Validation metrics for empirical Bayes models fit with progressively richer covariate and control-gene adjustment sets.

**Supplementary Figure S5.**
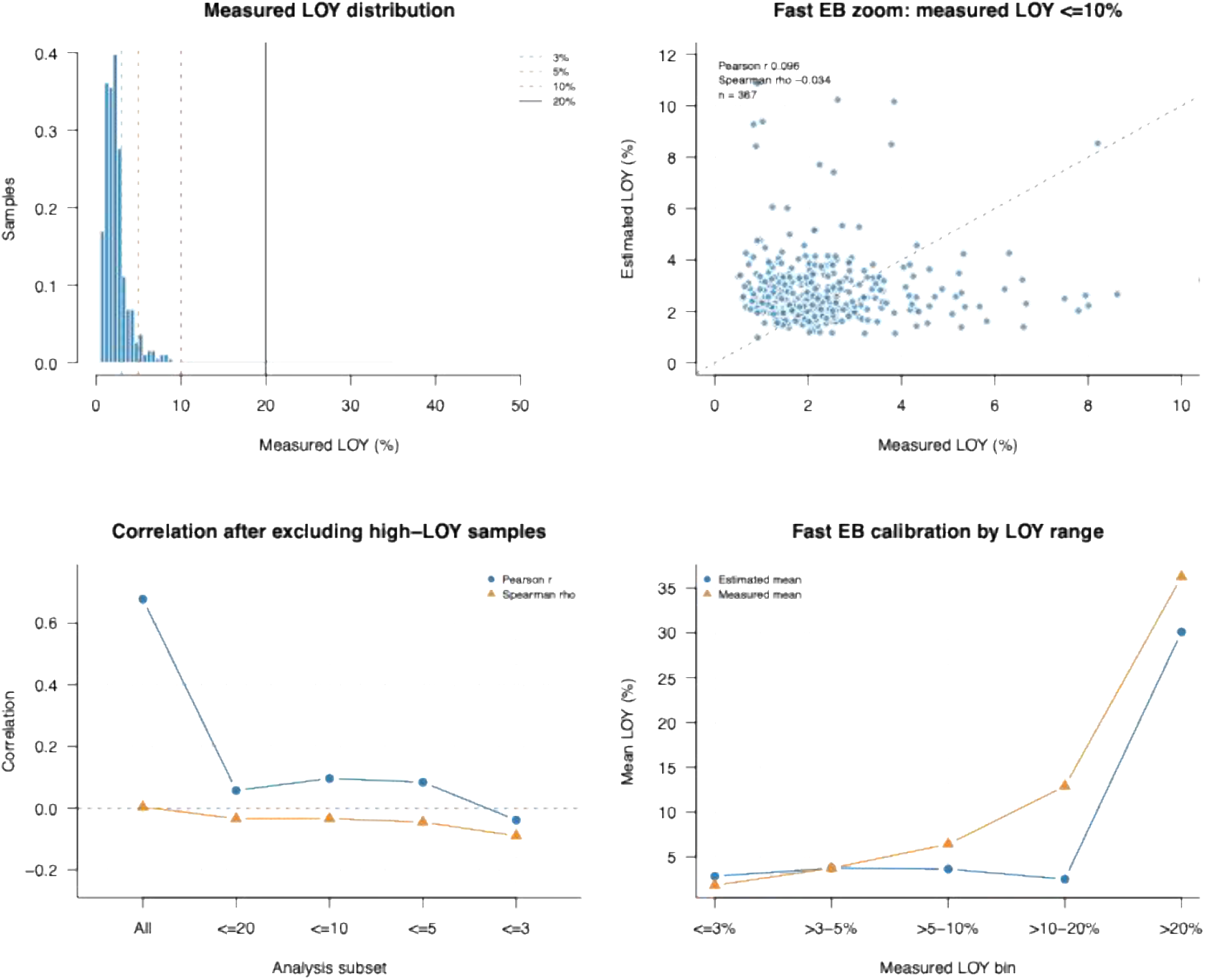
Low-LOY and high-outlier diagnostics. Distribution of measured LOY in the GTEx validation samples, Fast EB scatterplot restricted to measured LOY ≤10%, correlations after excluding high-LOY samples, and mean observed versus estimated LOY by measured LOY range. These diagnostics show that overall Pearson correlation is driven mainly by the small number of high-LOY samples, whereas monotonic ranking within the low-LOY range is limited.

## SUPPLEMENTARY TABLES

Supplementary Tables.xlsx includes the following supplementary tables:

- **Supplementary Table S1. Full bootstrap metric table**.
- **Supplementary Table S2. Hierarchical Bayesian posterior summaries**.
- **Supplementary Table S3a. GSE279480 sample manifest**.
- **Supplementary Table S3b. GSE279480 preprocessing summary**.
- **Supplementary Table S4. Gene-level stimulation effects**.
- **Supplementary Table S5. Low LOY and outlier sensitivity metrics**.
- **Supplementary Table S6. LOY range stratified calibration**.

